# Kinetics of reformation of the S_0_ state capable of progressing to the S_1_ state after the O_2_ release by Photosystem II

**DOI:** 10.1101/2024.09.10.612210

**Authors:** Alain Boussac, Julien Sellés, Miwa Sugiura

## Abstract

The active site for water oxidation in Photosystem II (PSII) comprises a Mn_4_CaO_5_ cluster adjacent to a redox-active tyrosine residue (Tyr_Z_). During the water-splitting process, the enzyme transitions through five sequential oxidation states (S_0_ to S_4_), with O_2_ evolution occurring during the S_3_Tyr_Z_^•^ to S_0_Tyr_Z_ transition. Chloride also plays a role in this mechanism. Using PSII from *Thermosynechococcus vestitus*, where Ca and Cl were replaced with Sr and Br to slow the S_3_Tyr_Z_^•^ to S_0_Tyr_Z_ + O_2_ transition (*t*_*1/2*_ ∼ 5 ms at room temperature), it was observed that the recovery of a S_0_ state, defined as the state able to progress to S_1_, exhibits similar kinetics (*t*_*1/2*_ ∼ 5 ms). This suggests that in CaCl-PSII, the reformation of the functional S_0_ state directly follows the S_3_Tyr_Z_^•^ to S_0_Tyr_Z_ + O_2_ transition, with no additional delay required for the insertion of a new substrate water molecule (O5) and associated protons.

## Introduction

Photosystem II (PSII) in plants, cyanobacteria, and algae by splitting water is at the origin of the atmospheric dioxygen and of the electrons used for making carbohydrates, *e*.*g*., [1] for a review. The PSII of *Thermosynechococcus vestitus* (formely *T. elongatus*), the thermophilic cyanobacterium used in this work, consists of 17 trans-membrane proteins and 3 extrinsic membrane proteins [2]. These proteins bind the cofactors involved in the trapping of light and the electron transfer reactions that are: 35 chlorophylls *a* (Chl-*a*), 2 pheophytins (Phe), 1 membrane b-type cytochrome, 1 PSII bound c-type cytochrome (replaced by a 17 kDa protein in plant PSII), 1 non-heme iron, 2 plastoquinones 9 (Q_A_ and Q_B_), the Mn_4_CaO_5_ cluster, 2 Cl^-^, 12 carotenoids and 25 lipids [2].

After the absorption of a photon by one of the 35 chlorophylls of the antenna the excitation energy is transferred to the Chl_D1_, forming *Chl_D1_. Chl_D1_ belongs to the photochemical trap that consists of the four Chls; P_D1_, P_D2_, Chl_D1_, Chl_D2_. These 4 Chls, together with the 2 Phe molecules, Phe_D1_ and Phe_D2_, constitute the reaction center of PSII. A few picoseconds after the formation of the excited *Chl_D1_, a charge separation occurs resulting ultimately in the formation of the Chl _D1_^+^ Phe _D1_^-^ and then P _D1_^+^ Phe D1^-^ radical pair states, *e*.*g*. [3-5], and *e*.*g*. [6,7] for recent theoretical works on this subject. After the charge separation, P_D1_^+^ oxidizes Tyr, the Tyr161 of the D1 polypeptide, and the radical Tyr_Z_^•^ is then reduced by the Mn CaO cluster. The electron on Phe _D1_^-^ is transferred to Q_A_, the primary quinone electron acceptor, and then to Q_A_, the second quinone electron acceptor. Whereas Q_A_ can be only singly reduced under normal conditions, Q_B_ is doubly reduced and protonated before to leave its site and replaced by an oxidized Q_B_ from the membrane pool, *e*.*g*. [8-11] and references therein.

The Mn_4_CaO_5_ cluster cycles through five redox states denoted S_n_, where n stands for the number of stored oxidizing equivalents. The S_1_-state is stable in the dark and therefore S_1_ is the preponderant state upon dark adaptation. When the S_4_-state is formed after the 3^rd^ flash of light given on dark-adapted PSII, two water molecules bound to the cluster are oxidized, O_2_ is released and the S_0_-state is reformed [12,13].

Thanks to the advent of serial femtosecond X-ray free electron laser crystallography (XFEL), the structure of the Mn_4_CaO_5_ cluster have been resolved in the dark-adapted state with the 4 Mn ions in a redox state as close as possible to that in the S_1_-state, *i*.*e*. Mn^III^ _2_Mn^IV^_2_[2,14-20]. The Mn_4_CaO_5_ structure resemble a distorted chair including a μ-oxo-bridged cuboidal Mn_3_O_4_Ca unit with a fourth Mn attached to this core structure *via* two μ-oxo bridges involving the two oxygen’s O_4_ and O_5_. Important progresses have been recently done in the resolution of the crystal structures in the S_2_ and S_3_ states [14-20]. The structural changes identified in the S_1_ to S_2_ transition, and those more numerous in the S_2_ to S_3_ transition cannot be detailed in a few lines. Very briefly, these works show that, when compared to S_1_, the changes in S_2_ are minor and more or less correspond to those expected for the valence change of the dangler Mn4 from +III to +IV, *e*.*g*., [21]. Importantly, water molecules in the “O1” and “O4” channels, defined as such because they start from the O1 and O4 oxygens of the cluster, appeared localized slightly differently in S_2_ than in S_1_ [17,20]. In contrast, in the S_2_ to S_3_ transition, major structural changes have been detected together with the insertion of a 6^th^ oxygen, originally a water molecule close to Ca and labelled O6 or Ox, finally bridging Mn1 and Ca [15,18-20]. The insertion of this 6^th^ oxygen was shown to be complete in 400 µs, and to occur simultaneously with the oxidation of Mn1 from +III to +IV [19,20]. This newly bound oxygen on Mn1 is supposed to correspond to the second water substrate molecule and is at a short distance of the bridging oxygen O5 supposed to be the first, and slowly exchangeable, bound water substrate molecule, *e*.*g*., [15,16,18,20,22]. An important movement of the Glu189 residue would allow its carboxylate chain to make a hydrogen bond with the protonated form of this 6^th^ oxygen in S_3_ [17,20].

There is a model, increasingly shared, in which after the 3^rd^ flash O_2_ is formed by a reaction between O5 and O6, even if the details including the way used by the released protons, are still being debated, *e*.*g*., [19,20,22-29].

Reformation of the S_0_ state after O_2_ evolution is less studied. This is partly explained by the fact that after each flash at least 10% of the PSII do not progress to the next S_n_ state. This implies that after 3 flashes the mix of the S_n_ states, *e*.*g*., [30], strongly complicates the interpretation of the data. The “S_4_” to S_0_ transition corresponds to the reduction of the Mn ions of the cluster, to the insertion to one of the two water substrate molecules and to some reprotonations [23,31]. Two to four ms after the 3^rd^ flash, the structure solved by XFEL of the Mn_4_CaO_5_ cluster and around, appeared to be the final one. Therefore, the S_0_ state appears to be reformed in all the centers in 4 ms [19], a time corresponding to the release of O_2_ in 100% of the centers [32,33]. It is however questionable whether this S_0_ state is fully functional and capable of being oxidized to S_1_, see [31,34] for recent computational works on this issue. Here, for an experimental kinetic approach, the advancement of the S_n_ state cycle was followed by measuring the ΔI/I induced at 292 nm by each flash of a sequence [33,35], and in which an (additional) extra saturating flash was given at varied time (Δt) after the third one, *e*.*g*. [36]. For the too short Δt values, the centers are expected to be unable to progress to the next S_n_ state, and the flash following this Δt should have no effect on the period 4 oscillation. When the Δt increases, a shift in oscillations is expected, as the additional flash has more and more the effect of a fourth flash, which gives the kinetics for the reformation of a functional S_0_ state.

## Materials and Methods

Culture of *T. vestitus* WT3* cells in the presence of Sr^2+^ and Br^-^, instead of Ca^2+^ and Cl^-^, was done as previously described [33]. Purification of the SrBr-PSII was done with the same protocol described in [37].

Time-resolved absorption changes measurements were performed with a lab-built spectrophotometer [38] slightly modified as detailed in [37]. Absorption changes were sampled at discrete times after the actinic flash by short weak measuring flashes. These measuring flashes were provided by an optical parametric oscillator (Horizon OPO, Amplitude Technologies) pumped by a frequency tripled Nd:YAG laser (Surelite II, Amplitude Technologies), producing monochromatic flashes (355 nm, 2 nm full-width at half-maximum) with a duration of 5 ns. Actinic flashes, for all the samples studied, were provided by a second Nd:YAG laser (Surelite II, Amplitude Technologies) at 532 nm, which pumped an optical parametric oscillator (Surelite OPO plus) producing monochromatic saturating flashes at 695 nm with the same pulse-length. The time delay between the laser delivering the actinic flashes and the laser delivering the detector flashes was controlled by a digital delay/pulse generator (DG645, jitter of 1 ps, Stanford Research). The path-length of the cuvette was 2.5 mm. The analytical flash was split into two parts using an optical guide composed by a bunch of randomized optical fibers. One part is connected to the reference cuvette for the measurement of the I_0_ and the other part is connected to the second cuvette, for the measurement of I, and to which is also connected the optic fiber from the actinic flash. The ΔI/I is equivalent to the (I-I_0_)/I value.

In this work, the absorption changes have been measured at 292 nm. This wavelength corresponds to an isosbestic point for Q_A_^-^/Q_A_, and PPBQ^-^/PPBQH_2_, the added electron acceptor, and is in a spectral region where the absorption of the Mn_4_CaO_5_ cluster depends on the S_n_-states which allow us to detect oscillations with a period four [35].

Once purified, the Ca and Sr do not exchange in the PSII [33] in contrast to the chloride, which can be replaced by bromide or iodide [39]. Therefore, for the ΔI/I measurements the samples were diluted in 1 M betaine, 15 mM CaBr_2_, 15 mM MgBr_2_, and 40 mM MES (pH 6.5). PSII samples were then dark-adapted for ∼ 1 h at room temperature (20–22 °C) before the addition of 0.1 mM phenyl *p*–benzoquinone (PPBQ) dissolved in dimethyl sulfoxide. The chlorophyll concentration of all the samples was ∼ 25 μg of Chl mL^-1^.

## Results and discussion

The constraint for measuring the “reopening” of the electron donor side of PSII after the 3^rd^ actinic flash given to dark-adapted PSII is that the kinetics for the reoxidation of Q_A_^-^ must be faster than the S_3_Tyr_Z_^•^ to S_0_ +O_2_ step. Since the electron transfer from Q_A_^-^ to Q_B_ occurs with a *t*_1/2_ value between 500 µs and ∼ 1-2 ms, *e*.*g*. [40], these conditions can only be fulfilled in PSII in which the Ca and Cl have been replaced by Sr and Br [33], and in which the *t*_1/2_ for the S_3_ Tyr_Z_^•^ to S_0_ +O_2_ step was slowed down to ∼ 5-7 ms.

Fig. 1 shows a control experiment in which the amplitude of absorption changes induced by each saturating flash, at 695 nm, of a series (200 ms apart) in the SrBr-PSII (with PPBQ present) used in this work was measured. The ΔI/I were measured by using a weak measuring flash at 292 nm, given 100 ms after each flash of the sequence. The oscillating pattern with a period of four [35] is similar to that already reported, and described in detail, in the SrBr-PSII [33].

**Figure 1:**
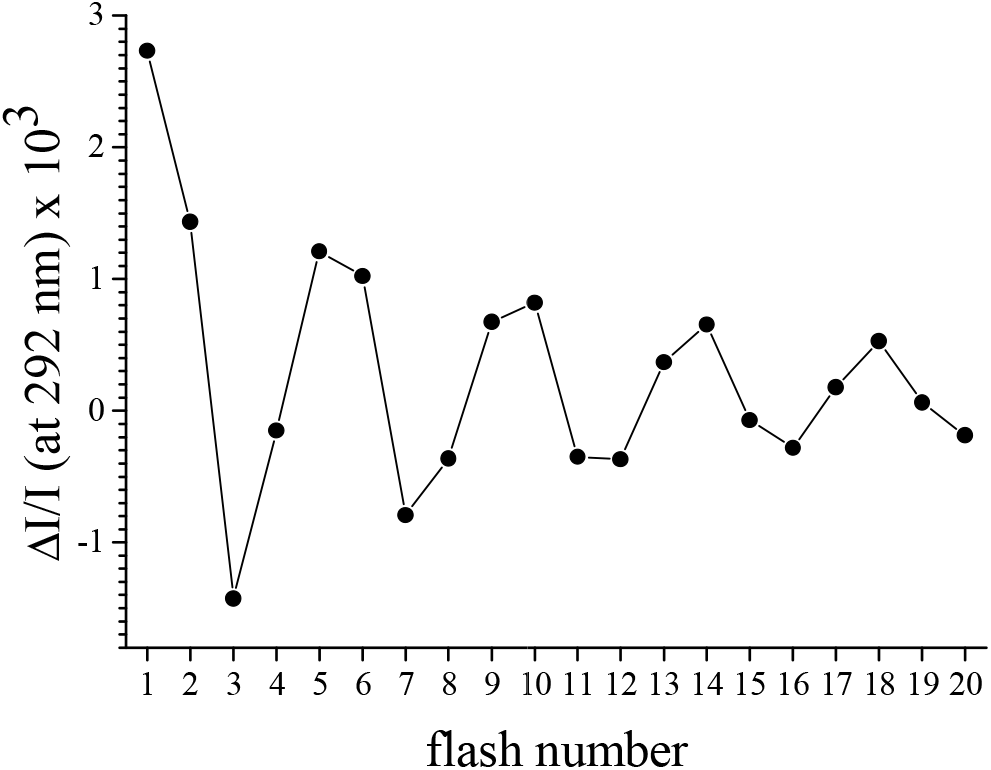
Sequence of the amplitude of the absorption changes at 292 nm. Measurements were done during a series of saturating flashes (spaced 200 ms apart) given on SrBr-PSII. The sample (Chl = 25 µg mL^-1^) was dark-adapted for 1 h at room temperature before the addition of 100 µM PPBQ (dissolved in dimethyl-sulfoxide). The term “flash number” refers to the number of actinic flashes applied to the dark-adapted sample. The signal (ΔI/I on the Y-axis) was measured using a weak flash at 292 nm, 100 ms after each actinic flash. The continuous line is drawn for joining the data points.

Fig. 2 shows the absorption changes, at 292 nm, from 10 μs (*i*.*e*. in the S_n_Tyr_Z_^•^state) to the ms time range (*i*.*e*. in the S_n+1_Tyr_Z_ state) after the first 3 flashes given to a dark-adapted SrBr-PSII. This experiment allows us to assess the kinetics of the S-state transition after the first flash (black), the second flash (red) and the third flash (blue). At 292 nm, the absorption changes associated with the S_2_Tyr_Z_^•^ → S_3_Tyr_Z_ transition on the second flash (red trace) are small and preclude a reliable kinetic analysis. As already observed [33], the kinetics of the absorption changes in the S_1_Tyr_Z_^•^ → S_2_Tyr_Z_ (black trace) with a *t*_1/2_ ∼ 200 µs is slightly slower in SrBr-PSII than in CaCl-PSII in which the *t*_1/2_ is ∼ 50 µs. In the S_3_Tyr_Z_^•^ to S_0_Tyr_Z_ transition (blue trace), two phases are observed [32,41,42]. The fast phase with a *t*_1/2_ ∼ 50 to 100 μs in *T. vestitus* [33], which corresponds to the release of the first proton in this transition before the reduction of Tyr_Z_^•^ by the Mn cluster, is seen as a lag phase at 292 nm [43]. The slow phase, seen here as an absorption decay with a *t*_1/2_ ∼ 5 ms corresponds to the S_0_ and O_2_ formations and to the release of an additional proton, *e*.*g*., [43,44].

**Figure 2:**
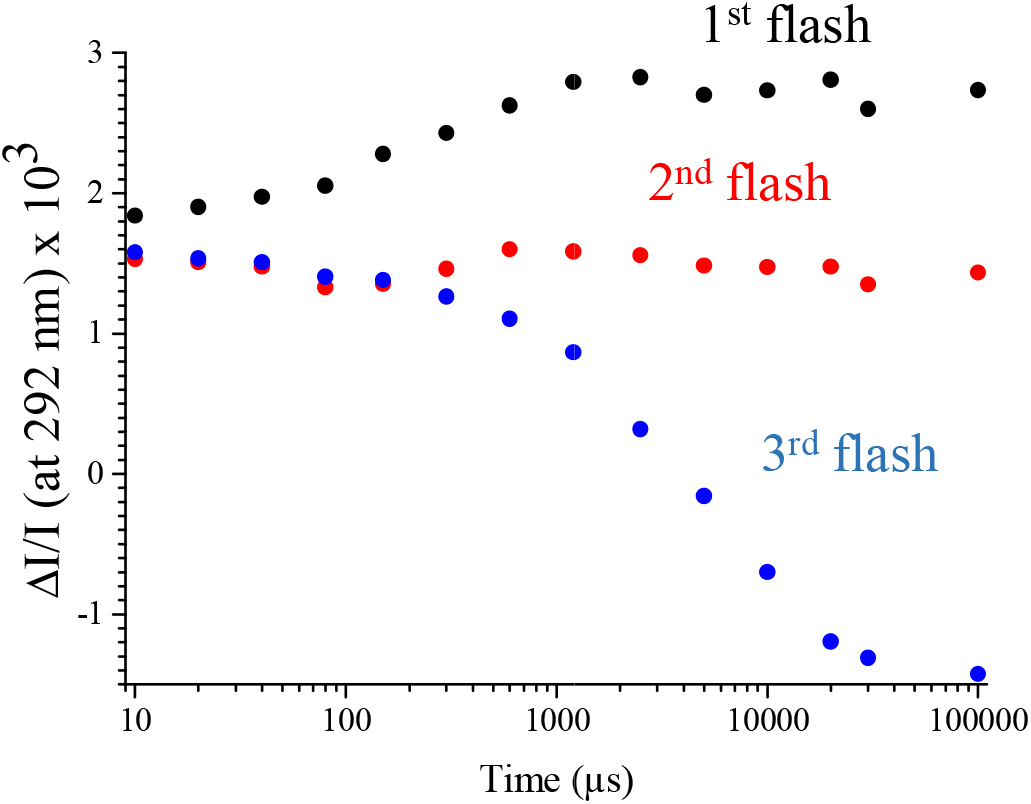
Kinetics of the absorption changes at 292 nm after the first flash (black points), the second flash (red points), and the third flash (blue points), given to dark-adapted SrBr-PSII. Other experimental conditions were similar to those in Fig. 1.

To ensure that the kinetics for the oxidation of Q_A_^-^ is indeed significantly faster than the S_3_Tyr_Z_^•^ to S_0_Tyr_Z_ transition, an experiment was carried out to estimate the reoxidation of Q_A_^-^ in the SrBr-PSII used here. Panel A in Fig. 3 shows the protocol used. It is based on the approach in [36]. In this experiment, the ΔI/I were recorded 100 ms after each flash of a sequence (green arrows) alternately without and with an extra saturating flash (red arrow) given Δt after the first flash. The sequence without the extra flash provides as many ΔI/I values corresponding to the first flash as sequences with the extra flash after the first flash, thus avoiding using the same data corresponding to the first flash for all Δt values.

**Figure 3:**
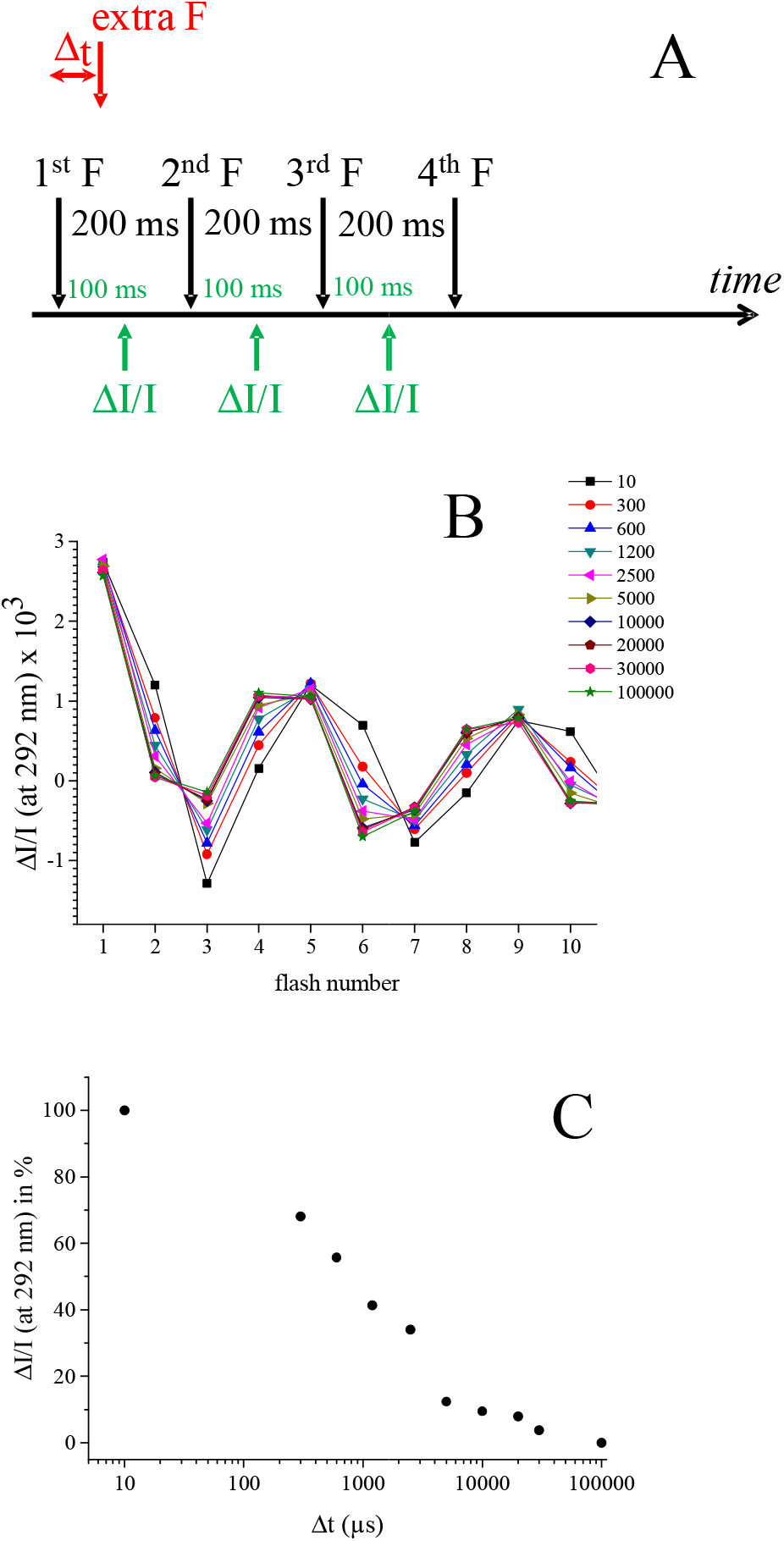
Panel A, schematic view of the protocol used. An extra actinic flash was given Δt after the 1^st^ actinic flash; Panel B, oscillations of the ΔI/I after each flash of a sequence *versus* the Δt between the first flash and the extra flash to SrBr-PSII; Panel C, Amplitude of the ΔI/I measured after the third flash *minus* the ΔI/I measured after the second flash (normalized to 100 % for Δt = 10 µs and to 0 % for Δt = 100 ms). Other experimental conditions were similar to those in Fig. 1.

Panel B in Fig. 3 shows the flash dependent ΔI/I values for Δt values ranging from 10 µs to several ms (the color code for the Δt in µs is on the right part of the Panel). When the extra flash was given 10 µs after the first flash the centers were still closed with both Tyr_Z_^•^ and Q_A_^-^ being largely present. Consequently for ΔT = 10 µs, the ΔI/I after the 3^rd^ flash is close to that after the 3^rd^ flash in Fig. 1 since the extra flash has no effect. For longer Δt values between the first flash and the extra flash in red in Panel A of Fig. 3 the reaction centers become more and more “reopen”. The reopening occurs firstly on the donor side and then, in a second phase, on the acceptor side, so that the flash labelled “2^nd^” flash on the X-axis of panel B in Fig. 3 corresponds actually to a 3^rd^ flash as defined in Fig. 1. The kinetics of the reopening of the centers after the first flash was estimated by plotting the ΔI/I measured after the 3^rd^ flash *minus* the ΔI/I measured after the 2^nd^ flash. Panel C in Fig. 3 shows this plot. Despite the limited number of data points, we can hypothesize that the decay observed in the 100 to 200 µs time range corresponds well to the S_1_Tyr_Z_^•^ → S_2_Tyr_Z_ kinetics. For its part, the slow phase with a *t*_1/2_ ∼ 1 ms fit well with the reoxidation of Q_A_^-^ [40,45]. Two comments can be made here: *i*) the fast phase is detected probably in centers in which Q_A_^-^ is oxidized by the non-heme iron with a kinetics much faster than the S_1_Tyr_Z_^•^ → S_2_Tyr_Z_ kinetics, and *ii*) the reoxidation of Q_A_^-^ (*t*_1/2_ ∼ 1 ms, [40,45]) occurs more rapidly than the S_3_Tyr_Z_^•^ to S_0_Tyr_Z_ transition (*t*_1/2_ ∼ 5 ms), [33] and Fig. 2. The *t*_1/2_ for the reoxidation of Q_A_^-^ by the oxidized non-heme iron has been estimated to be ∼ 55 µs [10]. This kinetics has been measured with DCMU bound, a condition known to increase the *E*_m_ of Q_A_ by ∼ 50 mV [46,47] and therefore to likely slow down the electron transfer from Q_A_^-^ to Fe^3+^. That means that the value of 55 µs is probably an upper limit in the absence of DCMU.

The conditions are now established for measuring the kinetics at which a functional S_0_ state is formed after 3 flashes and after O_2_ is evolved. Panel A in Fig. 4 shows the protocol used for that. In this experiment, the ΔI/I were also recorded 100 ms after each flash of a sequence (green arrows) alternately without and with an extra flash (red arrow) given Δt after the third flash. The sequence without the extra flash drawn in red in Panel A, provides as many ΔI/I values corresponding to the first 3 flashes as sequences with the extra flash, thus avoiding using the same data corresponding to these first 3 flashes for all Δt values.

**Figure 4:**
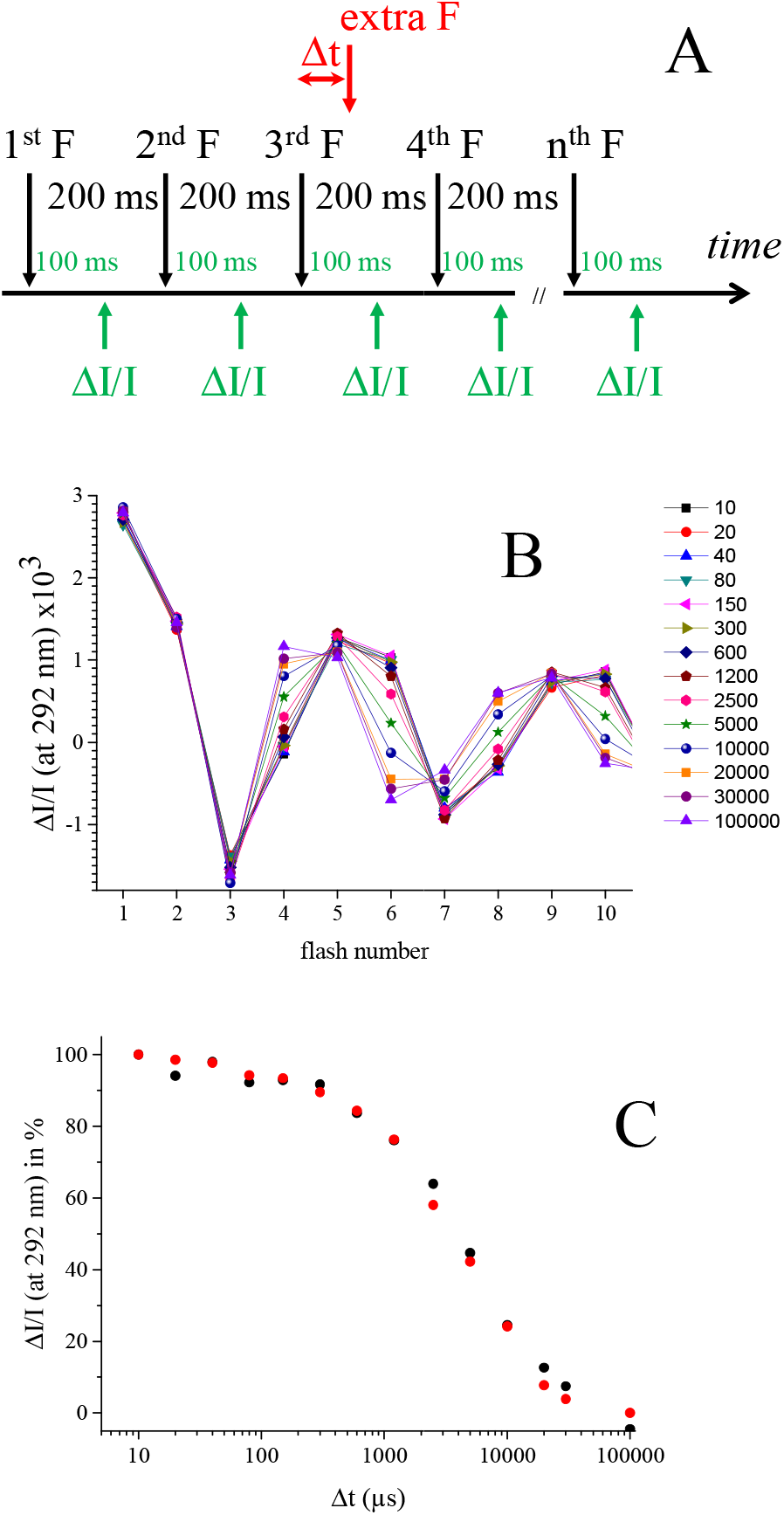
Panel A, schematic view of the protocol used when the extra actinic flash was given after the third actinic flash; Panel B, oscillations of the ΔI/I after each flash of the sequence *versus* the Δt between the third flash and the extra flash to SrBr-PSII; Panel C; The black points are the amplitude of the ΔI/I measured after the fourth flash *minus* the ΔI/I measured after the third flash (in Panel B, and the red data points are the ΔI/I after the third flash and shown in Fig. 2 The values are normalized to 100 % for Δt = 10 µs and to 0 % for Δt = 100 ms. Other experimental conditions were similar to those in Fig. 1.

Panel B in Fig. 4 shows the flash dependent ΔI/I values with Δt values ranging from 10 µs to several ms between the 3^rd^ flash and the extra flash (the color code for the Δt in µs is on the right part of the Panel). As for the experiment reported in Fig. 3, when the Δt was 10 µs, both Tyr_Z_^•^ and Q_A_^-^ remains largely present, so that the ΔI/I after the 4^th^ actinic flash is close to that after the 4^th^ actinic flash in Fig. 1. For longer Δt values the flash labelled “4^th^” flash on the X-axis of panel B in Fig. 4 has a ΔI/I value close to that after the 5^th^ flash as defined on the X-axis of Fig. 1.

The black data points in Panel C of Fig. 4 shows the amplitude of the ΔI/I measured after the fourth flash *minus* the ΔI/I measured after the third flash (normalized to 100% for Δt = 100 ms). We can apply here the same reasoning as for the experiment reported in Fig. 3. For the shortest Δt values, *e*.*g*., 10 µs between the 3^rd^ actinic flash and the extra flash drawn in red in Panel A, the centers are still in the S_3_Tyr_Z_^•^ state before the process resulting in the O_2_ formation and the before the release of protons begins. In other terms, it has no effect because the centers are closed. When the Δt increases, the proportion of S_0_ increases as the centers reopen. The red data points are a replot of the S_3_Tyr_Z_^•^ to S_0_Tyr_Z_ transition at 292 nm shown in Fig. 2 (after a normalization of the data from 100 % for the ΔI/I at t = 10 µs to 0 % for the ΔI/I at t = 100 ms). The resemblance between the 2 kinetics is remarkable. This resemblance shows that the kinetics of the formation of a S_0_-state capable of progressing to S_1_ follows the kinetics of O_2_ evolution without any apparent lag.

## Conclusion

After the third flash, the Mn_4_ cluster in the S_n_ state, called here S_0_, capable to progress to S_1_ with the same kinetics as the S_3_Tyr_Z_^•^ to S_0_ Tyr_Z_#x002B; O_2_ transition. This means than the rebinding of the water molecule to the cluster, *i*.*e*., the rebinding of O5, and all the proton bindings and movements [23,31,34] have a kinetics equal or faster than the regeneration of an S_0_ state capable of progressing to S_1_ after a fourth flash. This kinetics has been measured in SrBr-PSII in order the Q_A_^-^ reoxidation kinetics was not limiting. This imply that the kinetics of the recovery of a functional S_0_ corresponds to the situation in SrBr-PSII and the *t*_1/2_ ∼ 5 ms can only be considered as upper limit in CaCl-PSII. Our data do not allow us to kinetically identify all the intermediate states seen in the S_3_ Tyr_Z_ ^•^ to S_0_ Tyr_Z_+ O_2_, *e*.*g*., [19,48], however, from the strong similarity between the recovery of a functional S_0_ state and the S Tyr_Z_^•^ to S_3_ Tyr_Z_+ O_2_ transition in Panel C of Fig. 4, it sems likely that the reformation of a functional S_0_ state also follows the S_3_ Tyr_Z_ ^•^ to S_0_Tyr_Z_ + O_2_ transition in CaCl-PSII.

### Abbreviations

PSII: Photosystem II
Chl: chlorophyll
Chl_D1_/Chl_D2_: monomeric Chl on the D1 or D2 side, respectively
MES: 2-(*N*-morpholino) ethanesulfonic acid
P_680_: primary electron donor
P_D1_ and P_D2_: individual Chl on the D1 or D2 side, respectively, which constitute a pair of Chl with partially overlapping aromatic rings
Phe_D1_ and Phe_D2_: pheophytin on the D1 or D2 side, respectively
PPBQ: phenyl *p*–benzoquinone
Q_A_: primary quinone acceptor
Q_B_: secondary quinone acceptor
Tyr_Z_: the tyrosine 161 of D1 acting as the electron donor to P_680_
WT*3: *T. elongatus* mutant strain deleted *psbA*_*1*_ and *psbA*_*2*_ genes and with a His-tag on the carboxy terminus of CP43.
DCMU: 3-(3,4-dichlorophényl)-1,1-diméthyl-urée
XFEL: serial femtosecond X-ray free electron laser crystallography.

## Acknowledgements

This study was supported by i) the JSPS-KAKENHI Grant 21H02447 (MS), (ii) the Labex Dynamo ANR-11-LABX-0011-01 (JS).

